# Potential of Synthetic Aperture Radar Sentinel-1 time series for the monitoring of phenological cycles in a deciduous forest

**DOI:** 10.1101/2021.02.04.429811

**Authors:** Kamel Soudani, Nicolas Delpierre, Daniel Berveiller, Gabriel Hmimina, Gaëlle Vincent, Alexandre Morfin, Éric Dufrêne

**Affiliations:** Université Paris-Saclay, CNRS, AgroParisTech, Ecologie Systématique et Evolution, 91405, Orsay, France.; Institut Universitaire de France (IUF).; Laboratoire de Météorologie Dynamique, IPSL, CNRS/UPMC, Paris, France.

**Keywords:** Phenology, Forest, Sentinel 1, VV/VH, NDVI, LAI

## Abstract

Annual time-series of the two satellites C-band SAR (Synthetic Aperture Radar) Sentinel-1 A and B data over five years were used to characterize the phenological cycle of a temperate deciduous forest. Six phenological markers of the start, middle and end of budburst and leaf expansion stage in spring and the leaf senescence in autumn were extracted from time-series of the ratio (VV/VH) of backscattering at co-polarization VV (vertical-vertical) and at cross polarization VH (vertical-horizontal). These markers were compared to field phenological observations, and to phenological dates derived from various proxies (Normalized Difference Vegetation Index NDVI time-series from Sentinel-2 A and B images, *in situ* NDVI measurements, Leaf Area Index LAI and litterfall temporal dynamics). We observe a decrease in the backscattering coefficient (σ^0^) at VH cross polarization during the leaf development and expansion phase in spring and an increase during the senescence phase, contrary to what is usually observed on various types of crops. In vertical polarization, σ^0^VV shows very little variation throughout the year. S-1 time series of VV/VH ratio provides a good description of the seasonal vegetation cycle allowing the estimation of spring and autumn phenological markers. Estimates provided by VV/VH of budburst dates differ by approximately 8 days on average from phenological observations. During senescence phase, estimates are positively shifted (later) and deviate by about 20 days from phenological observations of leaf senescence while the differences are of the order of 2 to 4 days between the phenological observations and estimates based on *in situ* NDVI and LAI time-series, respectively. A deviation of about 7 days, comparable to that observed during budburst, is obtained between the estimates of senescence from S-1 and those determined from the in situ monitoring of litterfall. While in spring, leaf emergence and expansion described by LAI or NDVI explains the increase of VV/VH (or the decrease of σ^0^VH), during senescence, S-1 VV/VH is decorrelated from LAI or NDVI and is better explained by litterfall temporal dynamics. This behavior resulted in a hysteresis phenomenon observed on the relationships between VV/VH and NDVI or LAI. For the same LAI or NDVI, the response of VV/VH is different depending on the phenological phase considered. This study shows the high potential offered by Sentinel-1 SAR C-band time series for the detection of forest phenology for the first time, thus overcoming the limitations caused by cloud cover in optical remote sensing of vegetation phenology.

**Highlights:**

- We study S-1 C-band dual polarized data potential to predict forest phenology
- Seasonal phenological transitions were accurately described by S-1 time-series
- Budburst and senescence dates from S-1 differ from direct observations by one week
- Time-series of S-1 VV/VH, NDVI, LAI and litterfall were also compared
- Relationships VV/VH vs NDVI and LAI show a hysteresis according to the season

## 1. Introduction

In forest ecosystems, the opening of buds (“budburst”) in spring and the coloration and fall of leaves (“leaf senescence”) mark the start and the end of the photosynthetically active period and therefore play a key role in their productivity and carbon storage activity (Richardson et al., 2010). Historically, the timings of these phenological events have been monitored through direct and periodic human observations of trees in the field. However, this method is time-consuming, laborious, non-standardized and subject to an observer effect (Schaber and Badeck, 2002). Alternative indirect field techniques (RGB camera, proximal remote sensing systems, micrometeorological radiation sensors, etc.) are also used to monitor the seasonal cycle of forest canopy (Soudani et al., 2020). However, phenological metrics derived from both direct field observations and indirect proximal techniques are spatially spare and fail to describe the spatial and temporal variability in phenology due to the vegetation and micro-climate diversity (Soudani et al., 2012). Field observations also target a limited set of species and may not always be representative at the ecosystem scale. Satellite based remote sensing, because of its great potential for spatial sampling, constitutes the main approach for estimating and mapping phenological metrics at local to regional scales (Reed et al., 2003). However, the assessment of this potential has been severely limited by the difficulty of linking satellite-derived phenological metrics to field phenological observations, mainly due to temporal and spatial scale mismatches (Fisher et al., 2006). A few years ago, constellations of identical satellites such as SPOT 6/7, Pleiades, and more particularly, Sentinel-2 A and B in the optical domain and Sentinel-1 A and B in the C-band microwave frequency (5.405 GHz, 5.6 cm) have been launched, with the aim of overcoming these scale-related limitations by allowing image acquisitions with both good temporal and spatial resolutions under the same viewing angles (Ose et al., 2016). S2-A and S2-B, launched in 2015 and in 2017, that occupy the same orbit but 180° apart from each other, provide a temporal resolution of 10 days each and around 5 days with S2-A and S2-B together, reduced to 2-3 days over mid-latitudes regions. Spatial resolution of S2 varies from 10 m to 60 m depending on the spectral band. SAR (Synthetic Aperture Radar) S-1A and S-1B, launched in 2015 and 2016 respectively, together offer a temporal resolution about 3 days at the equator, > 1 day at high latitude and about 2 days in Europe combining ascending and descending orbits at a spatial resolution of 10 m in Interferometric mode (ESA, Sentinel-1 user guide). Temporal resolution of S-2 and S-1 are therefore comparable to the occurrence of phenological observations collected in the field over forest ecosystems, usually once or twice a week, and the spatial resolutions of 10 m and 20 m are also comparable to the size of adult forest tree crowns.

Typically, vegetation phenological metrics are derived from the analysis of time-series of spectral vegetation indices (SVI) in the optical domain but SVI are subject to varying degrees of uncertainty due mainly to cloud cover and cloud shadow contamination, which either introduces random noise that is difficult to correct or makes the data totally unavailable (Hird and McDermid, 2009; White and Wulder, 2013). Therefore, the temporal resolution of the satellite-based optical sensors is nominal since the availability of data depends on sky conditions (Wang and Atkinson, 2018; Sudmanns et al., 2019). In comparison to optical remote sensing, the main advantage of SAR remote sensing is its ability to pass through clouds with negligible attenuation. Temporal resolution is therefore maintained from year to year or from one region to another regardless of cloud conditions.

While the potential of S-2 for estimating forest phenology has been evaluated in many studies (Lange et al., 2017; Vrieling et al., 2018; Kowalski et al., 2020; Bolton et al., 2020), little is known about the potential of SAR data in general and S-1 in particular. The potential of S-1 has previously been assessed to monitor phenology, productivity and cultural practices in crops and meadows (Vavlas et al., 2020; Song and Wang, 2019; Stendardi et al., 2019), but at the exception of the study by Rüetschi et al. (2018), to the best of our knowledge no other studies which have compared phenological estimates derived from S-1 with field phenological observations in deciduous forests. In Rüetschi et al. (2018), few field phenological observations were available and, as pointed out by the authors, the temporal resolution of used S-1 time-series (24 days) was not adequate for an accurate assessment. In another studies, Dostálová et al. (2016; 2018) and Frison et al. (2018) analyzed time-series of S-1 over deciduous and coniferous forest stands. However, these studies were limited to the analysis of the temporal patterns of the S-1 data and did not focus on their exploitation for the detection of phenological dates.

In this paper, our first objective is to investigate the potential of time series of S-1 A&B C-band dual-polarized (VV and VH) SAR images to describe the phenological patterns of a temperate deciduous forest, and to estimate the timings of the main spring and autumn phenological stages. To this aim, we compared S1-based spring and autumn phenological estimates to in situ phenological observations by human observers and to estimates from alternative in situ indirect approaches including daily time-series of proximal NDVI (normalized difference vegetation index), daily time-series of LAI (Leaf Area Index) estimated from continuous radiation measurements and time-series of NDVI derived from S-2 A&B images. During the autumnal phenological stage, we also compared S1-based phenological dates to temporal dynamics of litterfall monitored in the field. Finally, and as a secondary objective of this paper, we further analyzed the sudden and abrupt changes observed in backscattering coefficients in the light of continuous measurements of precipitation and soil water content at different depths.

## 2. Materials and Methods

### 2.1. Site description

The study site is the Fontainebleau-Barbeau forest station (48°28’26”N, 2°46’57”E), located 53 km southeast of Paris (Supplementary Figure S1). Briefly, Fontainebleau-Barbeau forest is mainly composed of sessile oak (*Quercus petraea* (Matt.) Liebl), with an understory of hornbeam (*Carpinus betulus* L.). The stand age is about 150 years and the dominant height is 27 m. The topography is flat, and the ground elevation is about 103 m a.s.l. On this site which belongs to the pan-European ICOS Ecosystem network (Integrated Carbon Observation System, ICOS code FR-Fon), a 35-m high tower has been installed in 2005, measuring energy and matter (CO_2_ and H_2_O) exchanges between the vegetation and the atmosphere using the eddy-covariance technique. More details can be found in Delpierre et al. (2016) and at http://www.barbeau.u-psud.fr/index-fr.html.

### 2.2. Data

#### 2.2.1. SAR Sentinel-1 C-band time-series

Time-series of SAR Sentinel-1 (A&B) backscattering coefficient (σ^0^) at VH and VV polarization were composed using the Google earth engine (GEE) cloud. A total of 470 dual polarized (VV and VH) images covering the period from 01/01/2015 to 31/12/2019 were used. 225 scenes are in ascending orbit and 245 in descending orbits. Before September 2016 (day of year 278), S-1 time series are composed of S-1A images only. The number of S-1 images used per year is 43 images in 2015, 74 in 2016, 119 in 2017, 117 in 2018 and 117 in 2019; thus an average of about 1 image every 8 days in 2015, 1 image every 5 days in 2016 and 1 image every 3 days in 2017/2018 and 2019. As mentioned above, the number of images available in 2015 is low since only S-1A was operational. Images are in GRD (Ground Range Detected) format, in interferometric wide swath mode (IW), calibrated and ortho-corrected using the Sentinel-1 Toolbox at 10-m spatial resolution. Viewing incidence angle is about 39.30°. Each image contains two layers of the backscattering coefficient σ_0_ in VV and VH dual polarization converted in decibels unit (dB). Time-series of σ_0_ were composed based on their means within a circular buffer of 50 m in radius, centered on the Fontainebleau-Barbeau flux tower (Fig. S1). The number of S-1 pixels within the buffer is about 78 pixels. To remove noise, time series of VV/VH ratio in dB unit, calculated as (σ^0^VV (dB) - σ^0^VH (dB)), were filtered using the Savitzky-Golay filtering method under the MATLAB software. The best filtering was obtained with the following parameters: 2nd polynomial order and 5 or 9 window length.

#### 2.2.2. Sentinel 2 A & 2B time-series

A total of 284 Sentinel-2 (A&B) images (193 S-2A and 91 S-2B) from the S-2 L1-C TOA reflectance product were processed under GEE to generate NDVI time series using bands 4 (red) and 8 (near infrared) over the period 2015-2019. The corresponding wavelengths are respectively 664.5 nm (S-2A) / 665 nm (S-2B) for the red band and 835.1 nm (S-2A) / 833 nm (S-2B) for the near infrared band. The NDVI was calculated per pixel and averaged over the same 50 m diameter buffer as for S-1 A&B. The spatial resolution of the S-2 red and near infrared bands is 10 m and the number of pixels within the circular buffer is the same as for S-1 (78 pixels). Pre-filtering of cloudy pixels was performed using the QA60 flag, bits 10 and 11, which provide information about clouds and cirrus at the 60 m pixel scale. Only cloudy and cirrus-free pixels within the buffer were used in the calculation of NDVI. The number of S-2 images per year is 4 images in 2015, 11 in 2016, 17 in 2017, 28 in 2018 and 31 in 2019. From June 2015 to March 2017, only S-2A was operational, explaining the low number of images available for these two years. The year 2015 was excluded from the analysis due to the insufficient number of images.

#### 2.2.3 In situ data

##### 2.2.3.1. Field phenological observations

A description of field phenological data is given in Denéchère et al. (2019), Delpierre et al. (2020) and Soudani et al. (2020). Briefly, phenological observations were conducted according to two complementary sampling protocols. The first protocol (hereafter *intensive*) protocol was conducted from 2015 to 2017 for both spring and autumn. In this protocol, temporal dynamics of spring (percentage of open buds) and autumn (percentage of colored and/or fallen leaves) phenological transitions were determined at tree level, on 30 to 66 trees from the early signs to the end of each phenological stage. The second protocol (hereafter *extensive*) protocol was conducted on 2018 and 2019, during the spring phase only. In this protocol, we determined the date of budburst visually at the whole plot level surrounding the flux tower on about 100 trees. We considered that a tree had reached budburst when 50% of its crown showed open buds. Budburst of the whole plot was reached when 50% of sampled trees have reached budburst. All observations were achieved using binoculars, on a bi-weekly basis during the budburst (BB-OBS) and weekly during the senescence (LS-OBS). Hence the uncertainties are 3.5 days for BB-OBS and 7 days for LS-OBS.

##### 2.2.3.2. Narrow-band NDVI data

The NDVI is calculated as follows: *NDVI = (NIR - R)/(NIR + R)*. R and NIR are radiances in the red and the near infrared bands, respectively. Radiances are measured using a laboratory made NDVI sensor. A description of this sensor and its use for estimating phenological metrics in various biomes is given in Soudani et al. (2012; 2020) and Hmimina et al. (2013). Briefly, the sensor is positioned at the top of the flux tower in the Fontainebleau-Barbeau forest, about 7 m above the canopy, pointing downwards and inclined about 20-30° from vertical and facing south to avoid the hot-spot effects in canopy reflectance when the viewing direction is collinear with the solar direction. The field of view of the sensor is 100° and the observed area is a few tens of m². Measurements are acquired continuously every half-hour. Noisy data, due mainly to rainfall and very low radiation conditions, were removed according to the procedure described in Soudani et al. (2012). Daily average of filtered NDVI data acquired between 10h and 14h (UT) is considered to minimize daily variations in solar angle.

##### 2.2.3.3. Leaf Area Index LAI

For the different years, the maximum of leaf area index reached during the summer is determined directly using the litter collection method. 20 litter traps of 0.5 m^2^ each were used according to the standard protocol adopted within the framework of the European ICOS Ecosystem network (Gielen et al. 2018). The litter collection during the autumn is carried out at a one-week time step, thus allowing, in addition to the determination of maximum LAI, the description of temporal dynamics of surface area of fallen leaves from the first fallen leaves until the end of the autumn season.

Continuous estimation of canopy LAI was also achieved by applying the Beer-Lambert law to continuous measurements of incoming and beneath canopy radiation in the PAR (Photosynthetically active radiation) spectral range at a half-hourly time step. Beneath canopy radiation is measured using 15 sensors, installed on the ground-area surrounding the flux tower to ensure a robust spatial sampling of the radiation transmitted through the canopy. Since the application of BL’s law requires the prior determination of the extinction coefficient K, the latter was estimated from average LAI maximum determined from the litter collection method and average transmitted PAR during the summer growing season. To describe the temporal dynamics of LAI, this coefficient was assumed to be constant throughout the year. More details about the LAI calculation from radiation measurements are given in Soudani et al. (2020).

##### 2.2.3.4. Precipitations and soil moisture content

Rainfall and soil water content were measured at a half-hourly resolution. Rainfall is automatically measured using a rain gauge installed at the top of the tower (Model R01 3029, PRECIS MECANIQUE SAS, France). Soil water content is automatically measured every half hour and every 10 cm from the topsoil to 150 cm below the soil surface using 4 probes installed in the vicinity of the tower (SENTEK, Enviroscan system).

### 2.3. Methods

#### 2.3.1. Extraction of phenological metrics in spring and autumn from time-series

Six phenological metrics were extracted from daily time-series of S-1 VV/VH, in situ NDVI, S-2 NDVI, LAI and field phenological observations collected according to the intensive protocol. Three phenological metrics were also extracted from litterfall temporal dynamics. The extraction is carried out according to Soudani et al. (2008). Briefly an asymmetric double sigmoidal function (ADS) was fitted to time-series according to the following equation:

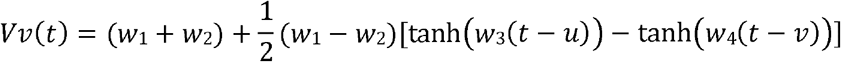

Vv (t) is the considered vegetation variable (VV/VH, NDVI from S-2 and in situ measurements, LAI, % of open buds, % of colored and/or fallen leaves). t is the time (day of year). tanh is the hyperbolic tangent and *w_1_*, *w_2_*, *w_3_*, *w_4_*, *u*, *v* are the fitted parameters. (*w*_1_+*w*_2_) is the Vv minimum in unleafy season. (*w*_1_-*w*_2_) is the total amplitude of the Vv seasonal cycle. The six phenological markers are named as follows: SOS, MOS and EOS for the start, middle, and end of season in spring and SOF, MOF and EOF for the start, middle and end of season in autumn, according Klosterman et al. (2014). The parameters were fitted by minimizing the sum of squares of differences between predicted and measured Vv. To constrain the fit at the end of the growing season, each year of VV/VH, NDVI and LAI data was extended to the winter of the following year. All dates were determined using the ADS function fitted to the data, except for SOF and EOF dates determined from litterfall temporal dynamics. For litterfall time-series, MOF is determined using the ADS function but SOF and EOF are determined directly from the data, due to the poor quality of the fit on both sides of ADS function. SOF and EOF are, respectively the dates corresponding to 10% and 90% of the total litterfall.

#### 2.3.2. Statistical analysis

The performance of VV/VH time-series to predict phenological events in spring and autumn was evaluated with respect to the field phenological observations using the mean bias error (MBE) and the mean absolute deviation (MAD) between estimated (*P_i_*) and observed dates (*O_i_*) for the different years (*N*), calculated as follows:

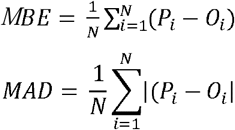

Statistical significance is determined at 5% probability level using one-tailed and two-tailed Student’s t-test for comparison of means. When linear regressions between different variables are investigated, R² is used to assess the strength of these relationships. Statistical analysis was done using the R software.

## 3. Results

### 3.1. SAR Sentinel-1 VV/VH time-series

S-1 VV/VH time-series are shown in Figure 1. σ^0^ in VV and VH polarization in ascending and descending orbits are shown in Figure S2, in supplementary material. Using S-1 original data (before filtering, Figure S2 for more details), the coefficient of backscattering σ^0^ is on average, over all years, about −8.54 dB ([−10.73, −5.07]) in VV polarization and about −14.40 dB ([−17.48, −10.45]) in VH. The difference between the two polarization is very highly significant (P < 0.001). Considering both ascending and descending orbits, differences in means are about 0.5 dB in both polarizations but are statistically significant. However, differences between means are not significant for VV/VH (P < 0.34). For this reason, S-1 data acquired at the two orbits are considered without distinction in the following.

**Figure 1:**
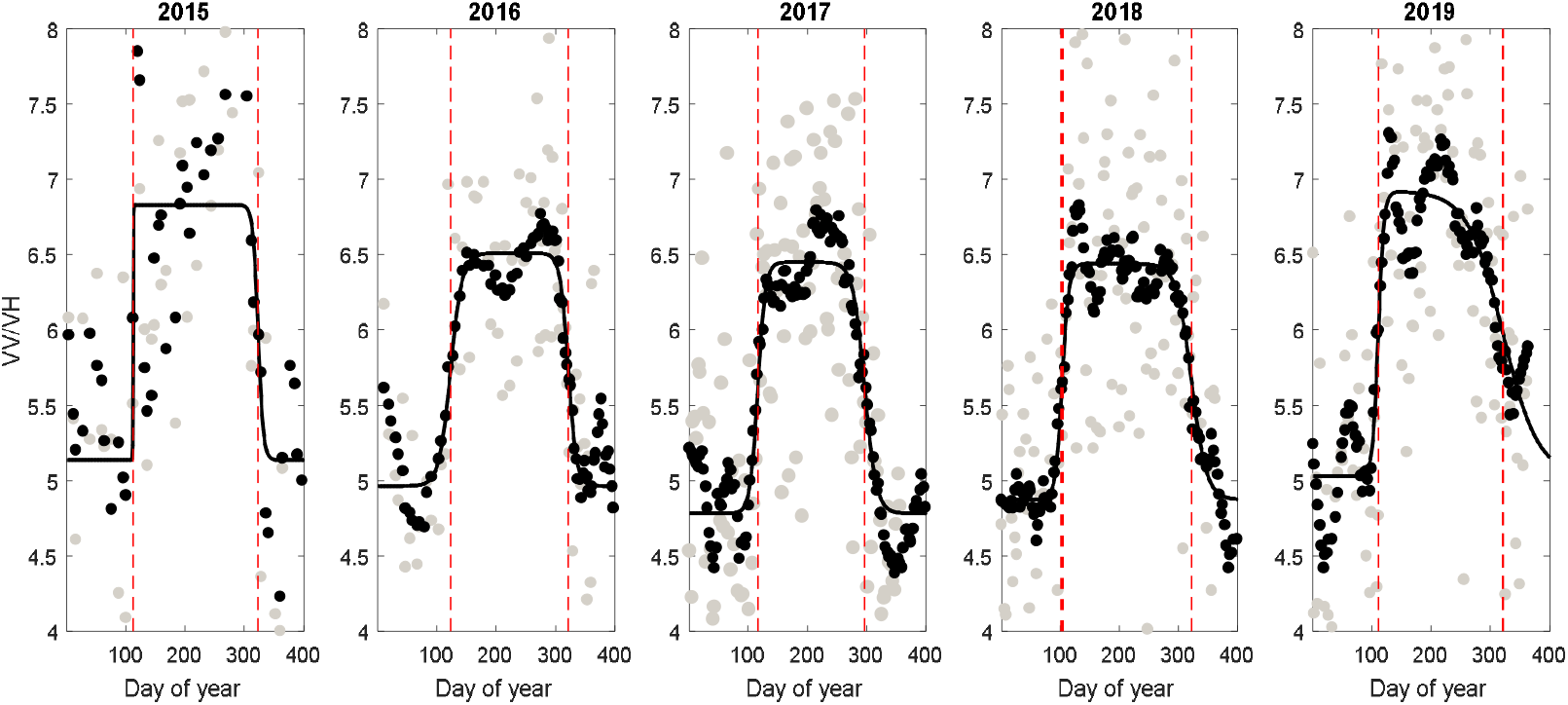
Sentinel-1 VV/VH time-series (dB): grey circle: original data; black circle: data smoothed using a Savitzky-Golay filter; continuous black curve: asymmetric double sigmoid function (ADS) fitted to VV/VH smoothed data. Red vertical bars: estimated phenological transition (MOS and MOF) dates from VV/VH time-series based on ADS function.

On the other hand, the analysis of the temporal dynamics of σ^0^VH and σ^0^VV shows intermittent sudden and abrupt changes. Weak relationships are observed between σ^0^VV, σ^0^VH, VV/VH and soil moisture between 0-10 cm over the year (R² 0.07; 0.26 and 0.148, respectively) but these relationships disappear when separating the summer leafy and winter unleafy seasons in the analysis (Figures S3 & S4). During the summer period, although weak, a significant positive relationship was obtained between rainfall and σ^0^VH (R² = 0.124, P < 0.008).

Figure 1 shows that S-1 VV/VH temporal patterns reproduce the forest phenology observed in deciduous forests with good accuracy. Four phases can be distinguished: a winter phase with low VV/VH values, a rapid transition phase during spring when VV/VH increases quickly, a plateau during the main growing season in summer and a fourth phase of VV/VH decline which coincides with the phase of leaf senescence and leaf fall. In comparison with the backscattering coefficients σ^0^VH or σ^0^VV, the VV/VH ratio is more dynamic and seems to be driven by VH more than VV polarization (Figure S2).

Indeed, the backscatter coefficient σ^0^VH decreases during spring phenological transitions, and remains relatively stable during the summer for which LAI remains stable and increases again during autumn phenological transitions (Figure S2). Between its winter maximum and summer minimum, σ^0^VH varies on average from −13.36 dB (standard deviation 0.871 dB) in winter [doy 330-98 next year] to −15.16 dB (standard deviation 0.851 dB) in summer [doy 128-270]. The decrease is of 1.80 dB, but σ^0^VH is highly significantly lower in summer than in winter (P <0.001, one-sided t-test). In vertical polarization, σ^0^VV decreases from −8.43 dB (0.732 dB) in winter to −8.51 (0.710 dB) in summer (Figure S2). The decrease is only about 0.15 dB but is statistically significant at 5% (P <0.048). In the VV/VH, driven by changes in σ^0^VH, the average increases from 4.93 dB (0.77 dB) in winter to 6.56 dB (0.79 dB) in summer. Also, VV/VH is very significantly higher in summer than in winter (P<0.001, one tailed t-test).

### 3.2. Phenological markers derived from SAR Sentinel-1 VV/VH time-series

The temporal dynamics of VV/VH strongly co-vary with the canopy phenological cycle, as assessed through various proxies (Figure 2). Summary statistics are given in Table 1 and estimated dates by year and for the six phenological metrics (SOS, MOS, EOS in spring and SOF, MOF and EOF in autumn) are given in Supplementary Material S5.

**Table 1:**
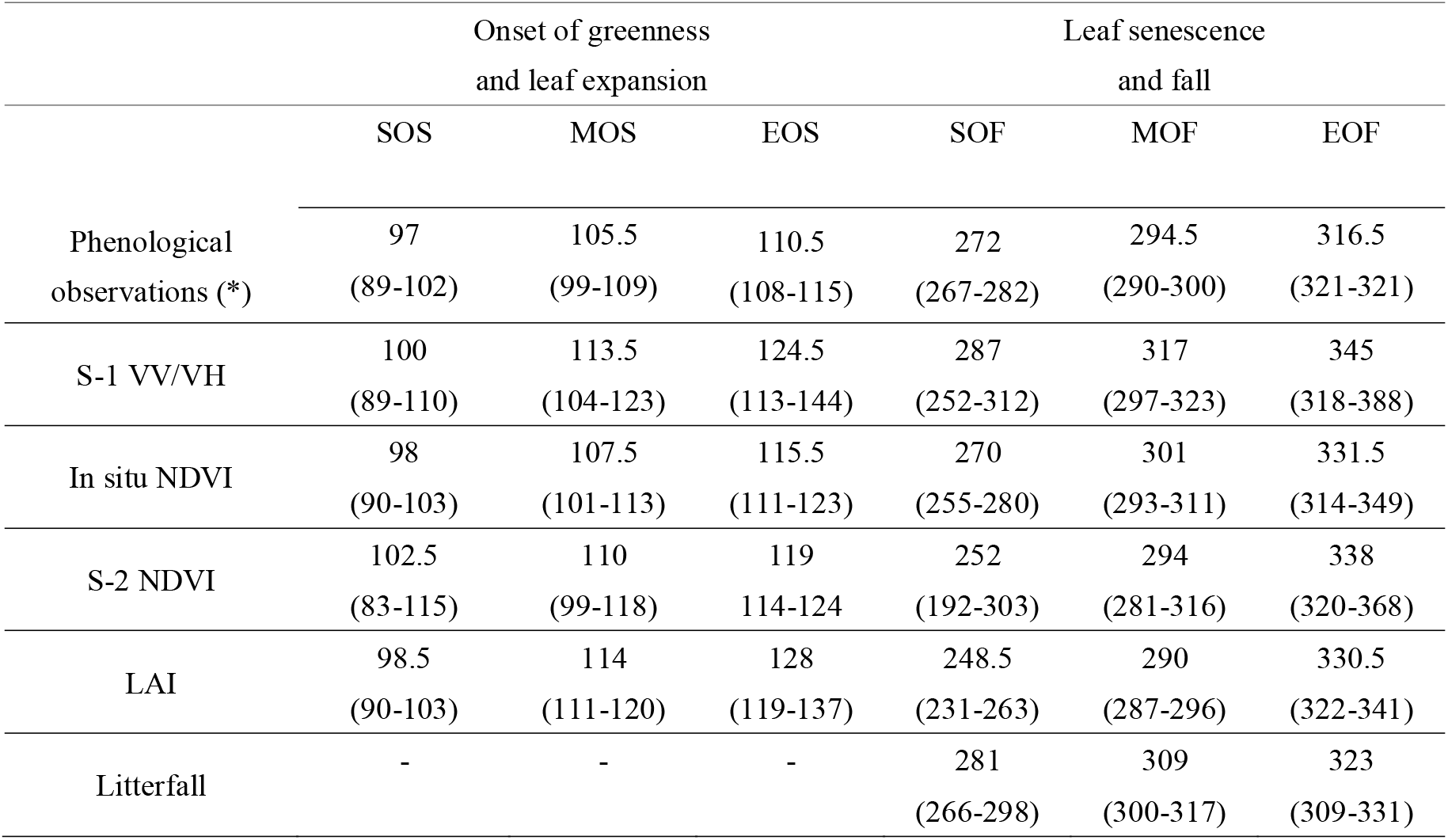
Summary statistics (mean and min-max) of observed and estimated phenological dates. SOS, MOS and EOS for the start, middle, and end of season in spring and SOF, MOF and EOF for the start, middle and end of season in autumn. (*) For field phenological observations, MOF is determined for the five years from the extensive protocol. SOS, EOS, SOF, MOF and EOF, are determined for years 2015 to 2017 using the intensive protocol.

**Figure 2:**
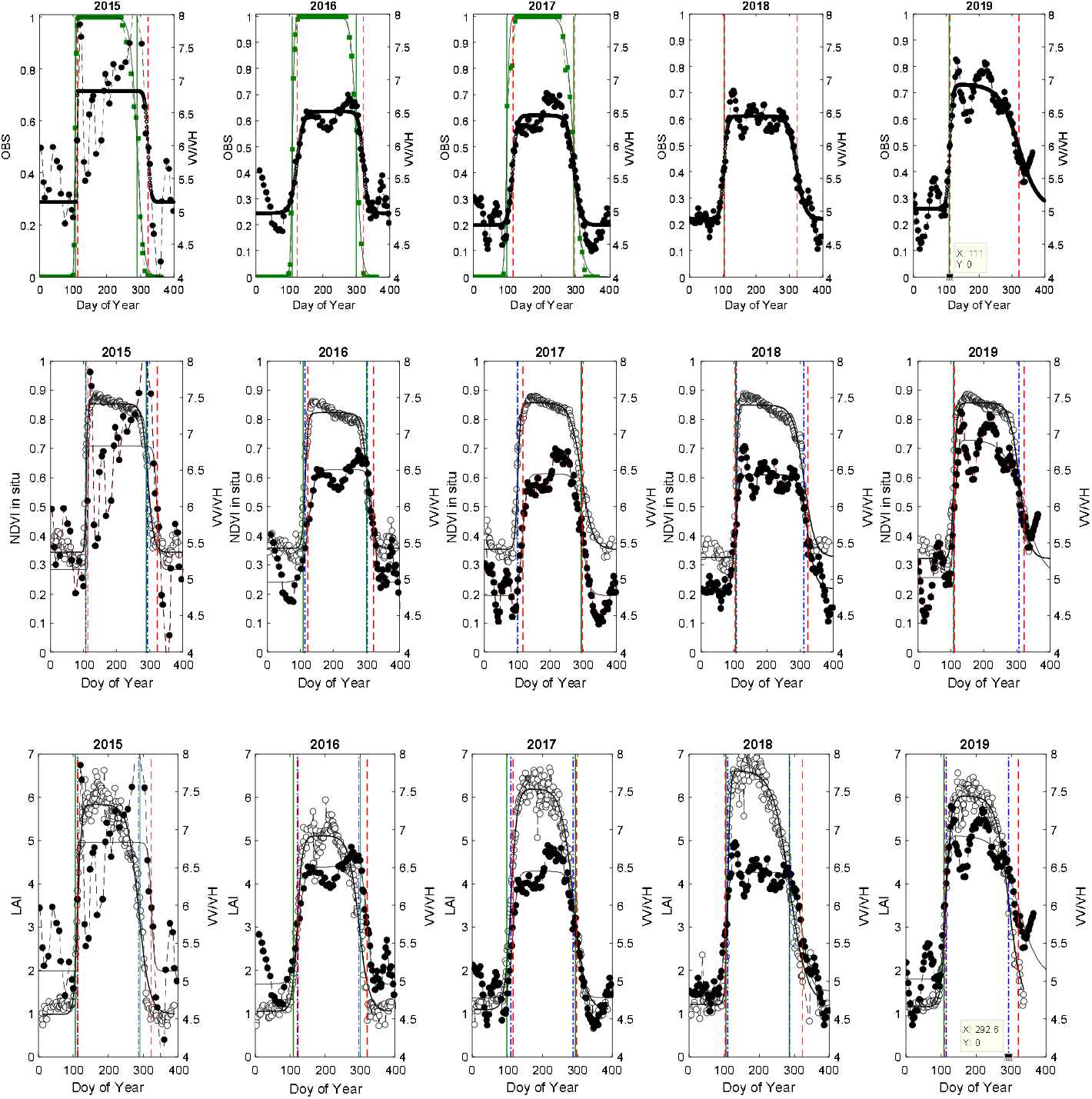
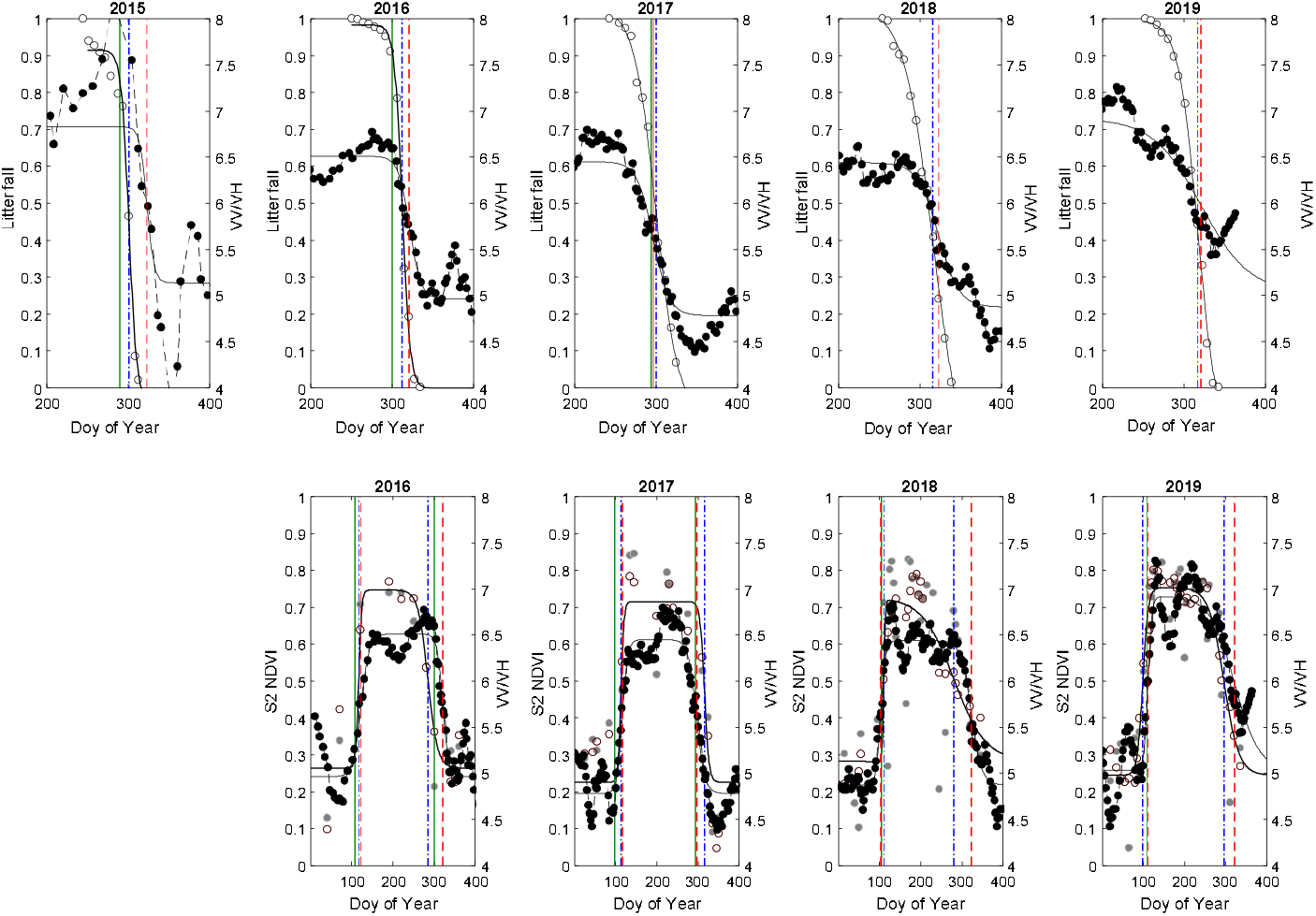
Sentinel-1 VV/VH time-series for all subplots (black filled circles – right axis). Empty circles – left axis: from top to bottom – time-series of field phenological observations (OBS from intensive protocol on 2015-2017), ground-based NDVI, Leaf Area Index (LAI), Litterfall in autumn and Sentinel 2 NDVI. Vertical bars: observed phenological transition dates in green (OBS from intensive and extensive protocols); estimated MOS and MOF phenological transition dates based on ADS function using VV/VH time-series in red and predicted phenological dates from in situ NDVI, LAI, S-2 NDVI (gray circle – removed using SG filter; empty circle – used) and litterfall in blue.

Figure 3 illustrates the interannual variations in estimated budburst (MOS) and senescence dates (MOF) for the different approaches.

**Figure 3:**
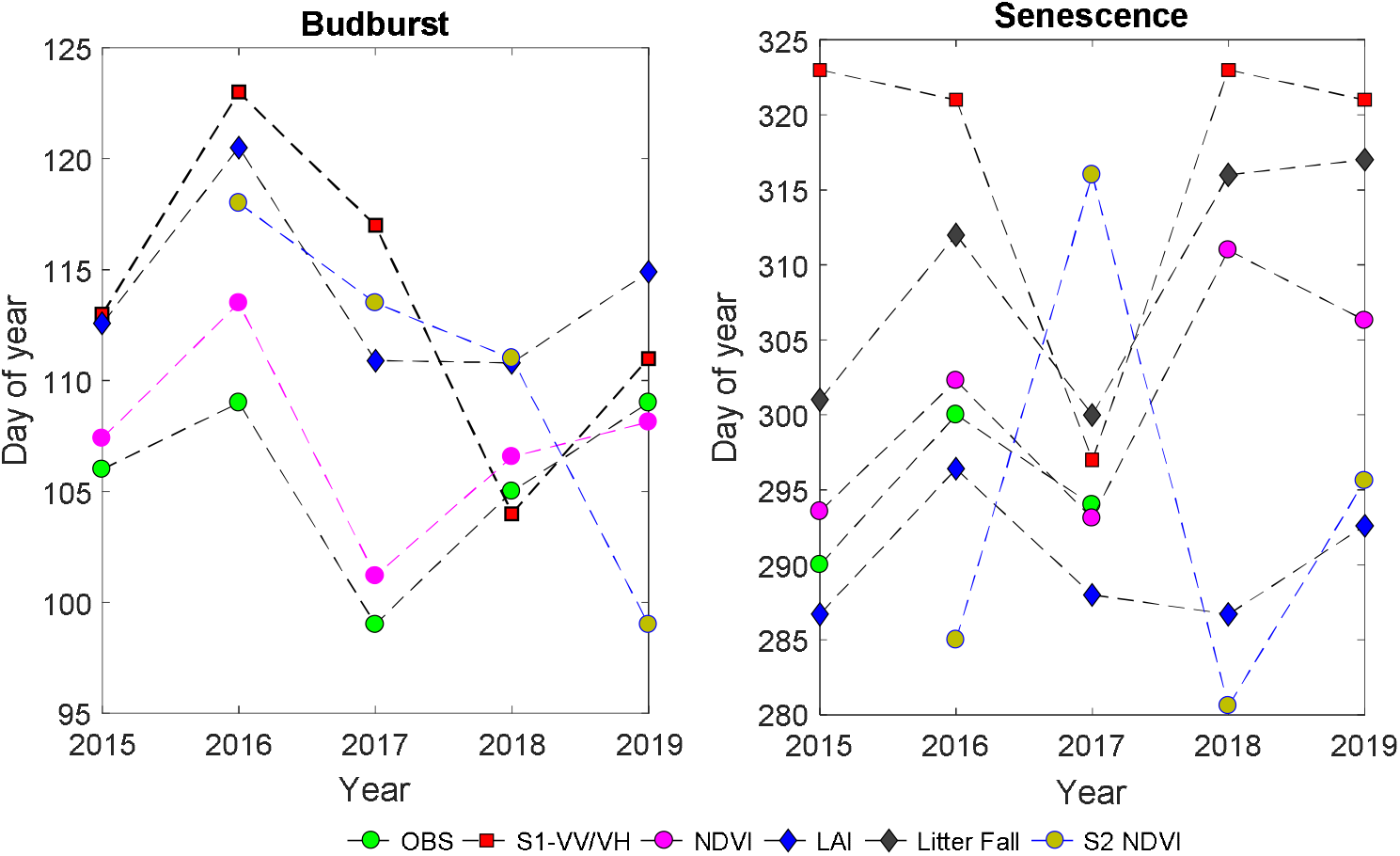
Average dates of budburst (a- left) and senescence (b-right) based on MOS and MOF, respectively.

During the spring transition phenological stage, and based on MOS criterion, VV/VH time-series lag about 8 days behind the observed budburst date (MBE and MAD, 8.5 and 8 days respectively; Figure 3a). This lag with observed budburst dates is similar to the lag of MOS retrieved from LAI time-series (MBE and MAD of about 8.5 days) and is slightly lower by about 1.5 days in comparison to the lag between MOS estimates retrieved from S-2 NDVI time-series (for S-2 NDVI, MBE and MAD of about 10 days and 5 days, respectively). The best estimates are obtained using in situ NDVI time-series for which MBE and MAD are about 2 days. During the autumnal phenological phase, for the three years for which continuous field observations are available, VV/VH time-series provide estimates which are approximately 20 days later than the observed dates (Figure 3b) while the differences are of the order of 2 to 4 days between the observations and estimates based on in situ NDVI and LAI, respectively. Interestingly, MOF derived from VV/VH differs by 7 days on average from MOF determined from litterfall time series. Between S-1 and S-2 based estimates, MBE and MAD are respectively of - 4 days (S2-based estimates later) and 7 days during the spring and −20 days (S-2 earlier) and 30 days in autumn (Figure 3b, Table S5).

These differences between VV/VH estimates and those obtained using field phenological observations and the other alternatives methods reflect different relationships between the temporal dynamics of the VV/VH signal and canopy properties. Figure 4 illustrates the relationships between VV/VH or LAI and in situ NDVI during four distinct phenological stages: winter dormancy period, spring leaf emergence and expansion stage, summer growth stage, and autumnal leaf senescence and fall. Four years, from 2016 to 2019, for which more S1 data are available and phenological estimates are more robust.

**Figure 4:**
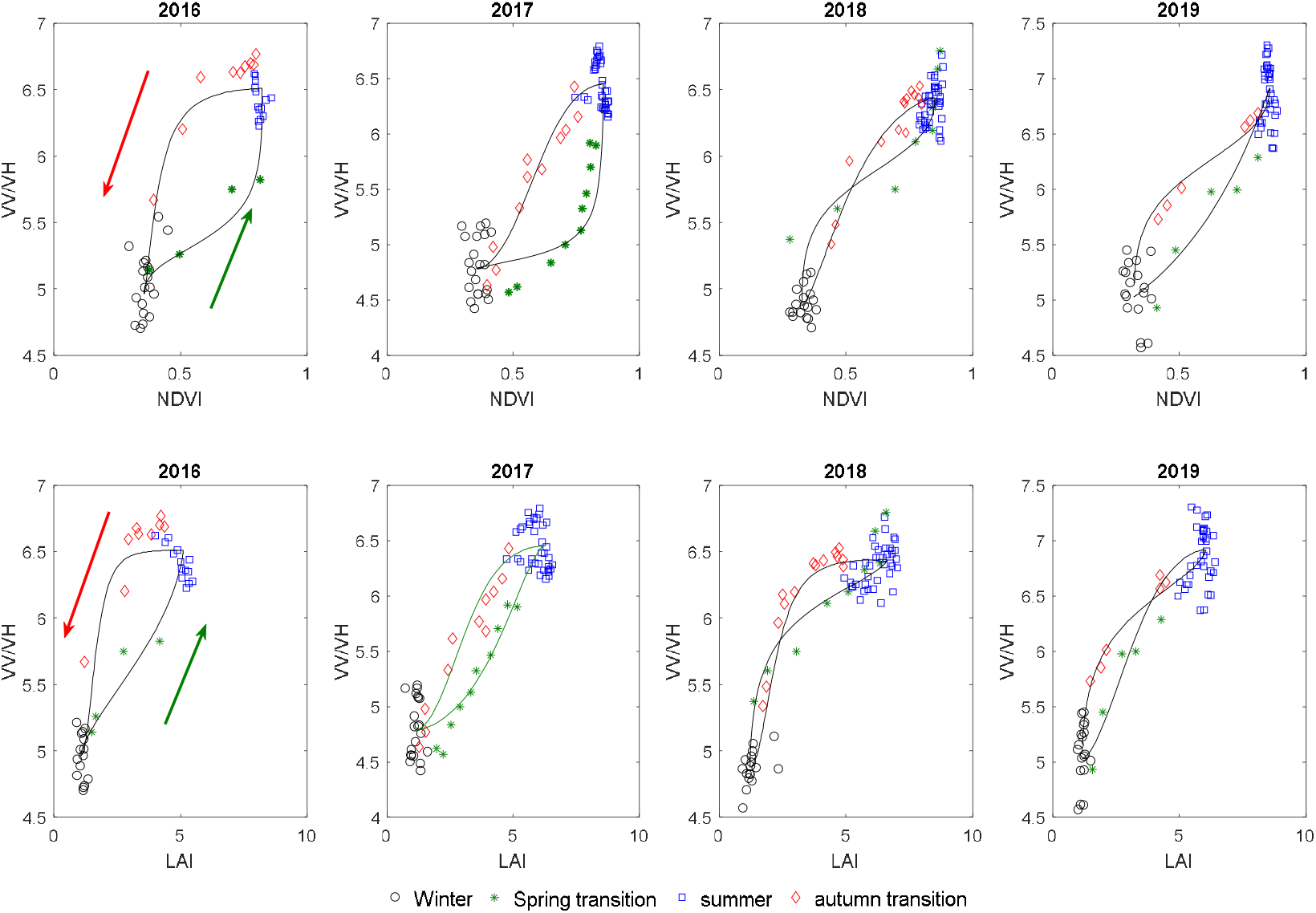
Relationships between VV/VH (dB) and in situ NDVI (top) and LAI (bottom) according to the phenological stage. Symbols are measurements: winter (day of year: 330-98 of the next year) in black circles; spring transition (day of year 98-128) in green stars; summer growing season (day of year 128-270) in blue square; autumn transition (day of year 270-330) in red diamond. Continuous curves are predicted values using the ASD function. Red and green arrows give the directions of VV/VH trajectories during the autumn (red) and spring (green) phenological stages, respectively.

Figure 4 shows a positive relationship between VV/VH and both NDVI and LAI. These relationships are not unique and depend on the phenological stage considered. A hysteresis phenomenon can be noted as the VV/VH trajectories are not identical during the increase and the decrease in canopy leaf area as described using LAI or NDVI, during spring and autumn, respectively. For the same NDVI (or LAI), VV/VH is higher during the fall phase than during the spring phase.

## 4. Discussion

Examination of the temporal dynamics of the VV/VH signal shows significant rapid variations that seem to be linked to rainy events (Figure 1, Figures S3 & 4). C-band sensitivity to precipitations intercepted and stored by the canopy during data acquisition has been shown in previous studies from measurements (Benninga et al., 2019; El Hajj et al., 2016; Proisy et al., 2000) and from simulations (Jong et al., 2016). As underlined above, the relationship between S-1 backscattering and soil water content is better in VH than in VV polarization and is only significant when both the unleafy and leafy seasons are combined (Figure S4). Therefore, this relationship seems to be caused more by the changes in canopy properties than by direct influences of soil moisture on backscatter coefficients. As also shown in Figure S3, σ^0^VH is more sensitive to the phenological cycle observed in deciduous forests while the temporal profile of σ^0^VV remains relatively stable throughout the year. σ^0^VH shows an average decrease of about 1.80 dB on average during summer in comparison to winter. This result agrees with the studies of Rüetschi et al. (2018), Dostálová et al. (2016; 2018) and Frison et al. (2018) in which they observe a lower backscattering coefficient in the S-1 VH polarization in summer than in winter in temperate deciduous forests. In Rüetschi et al. (2018), the differences observed on two oak stands and over two consecutive years ranged from 0.38 dB to 1.96 dB in VH polarization. In VV polarization, a lower average in summer than in winter was observed on a single oak stand. The difference was of 0.76 dB, but in general, there is no a well-established seasonal effect on σ^0^VV and the sign of the difference between summer and winter σ^0^VV varied. σ^0^VV was considered relatively stable throughout the year while σ^0^VH responds significantly to the seasonal vegetation cycle. In Rüetschi et al. (2018), the behavior of σ^0^VH was also verified over a whole region at the S-1pixel scale. Deciduous stands, composed mainly of oak and beech, showed most often a lower VH backscattering in summer than in winter. An opposite behavior was observed on coniferous stands, composed mainly of spruce, for which the σ^0^VH is higher in summer than in winter. Dostálová et al. (2016; 2018) also observed a clear decrease in S-1 VH signal during the spring and an increase during the senescence transition stage in the range of 0.5 to 2 dB in broadleaf forests of oak, beech, maple and birch trees. Similar results were obtained by Frison et al. (2018) over the whole Fontainebleau forest massif to which the Fontainebleau-Barbeau forest (our study site) belongs. The authors observed a decrease in σ^0^VH in spring from −12.5 to −15 dB, while σ^0^VV remains relatively constant.

Rüetschi et al. (2018) and Dostálová et al. (2016; 2018) explained the decrease in VH backscatter during the growing season, when leaf biomass is at its maximum, by a lower contribution of branches to backscattering, considered more reflective than leaves in C-band, and by a lower contribution of soil masked by foliage.

The sensitivity of σ^0^VH and VV/VH to phenology shown in our study (Figures 1 and S3) and in Rüetschi et al. (2018), Dostálová et al. (2016; 2018) and Frison et al. (2018), characterized by a decrease of the former and an increase of the latter during the summer, is different from what can be observed on various types of crops as shown in many studies (Veloso et al., 2017; Khabbazan et al., 2019; Stendardi et al., 2019; Dostálová et al., 2018) in which an increase in σ^0^VH signal and a decrease in VV/VH ratio when biomass increases in maize, soybean, sunflower, potatoes crops and in alpine meadows. In those studies, the increase of σ^0^VH backscattering was explained by the increase in vegetation volume scattering acting as the main backscatter.

As underlined above, VV/VH ratio is more dynamic than σ^0^VH and σ^0^VV and better describes the seasonal dynamics of the canopy (Figures 1 & S3). VV/VH is also known to be more stable and able to reduce the soil moisture and soil-vegetation interactions effects (Vreugdenhil et al., 2018; Veloso et al., 2017). As shown in Figure 2, S-1 VV/VH reproduces the annual cycle of phenological events observed in deciduous forests with varying degrees of fidelity.

The estimated phenological dates using MOS marker from VV/VH time-series are in agreement with the observed dates, with a positive bias of about 8 days during spring (Figure 3). A similar bias is obtained using the LAI time series. Indeed, during the spring transition phase (Figure 2), we observe that time-series of VV/VH overlap with those of LAI but deviate very significantly during the fall phase. During this phase, temporal patterns of VV/VH are positively shifted with respect to those of LAI. It can also be noted that temporal patterns of litterfall are also positively shifted in comparison to temporal patterns of LAI estimated from transmitted PAR. The average bias between estimates of the senescence date decreases from about 27 days between VV/VH and LAI to about 7 days between VV/VH and litterfall. During the senescence and compared to *in situ* and S-2 NDVI, VV/VH provides estimates that are generally later, between 4 and 30 days, with an average bias of 17 days compared to *in situ* NDVI and between −19 and 42 days and an average bias of 20 days with S-2 NDVI.

While during the spring phase, the patterns of VV/VH, LAI and NDVI from *in situ* and S-2 data, are relatively close, they deviate very strongly during the senescence phase causing large differences in the estimation of senescence dates (Figure 2). During this phase, VV/VH decay seems to more closely follow the temporal dynamics of litterfall than the LAI or NDVI decay. These results show that the relationships between VV/VH and LAI or NDVI are not stable but depend on the phenological stage considered. For the same LAI or NDVI, the VV/VH signal is generally weaker during the spring phenological stage than during senescence (Figure 4). This implies that the canopy characteristics that modulate these variables are different. During spring, the concomitance of VV/VH increase with the increase of LAI and to a lesser extent with NDVI reflects the sensitivity of VV/VH, and particularly VH, to leaf emergence and expansion. Leaf area development and growth act both in terms of number and geometrical properties of scattering elements composed of leaves and young twigs, changes in their dielectric properties especially through leaf water content and by masking the contribution of trunks, branches, and soil. It is very likely that this last factor is the most important because trees of the Fontainebleau-Barbeau forest are about 150 years old and the contribution of the woody parts is expected to play a major role in winter and to decrease as LAI increases during the spring. During this phenological stage, foliage development and expansion also modulate NDVI and LAI very strongly although they also depend on canopy structure and leaf structural and biochemical characteristics. During the senescence phase, the onset of VV/VH decline, estimated from the SOF criterion, is later than for NDVI and LAI and practically synchronous with litterfall. NDVI decreases before leaf fall due to leaf yellowing and browning that characterize autumn leaf senescence. During this period, in addition to the decrease in leaf chlorophyll content which is the main cause of the decrease in NDVI, leaf water content and leaf mass also decreased as shown in previous studies (Yang et al., 2016; Yang et al., 2017; Meerdink et al.,2016). LAI, calculated in this study from transmitted radiation, also decreases earlier, probably due to an increase of canopy transmissivity caused by a decrease in leaf chlorophyll content during this period. Therefore, VV/VH seems to depend much more on the amount of scattering elements (leaves, twigs, branches and trucks) which is better described by litterfall than by NDVI or LAI.

While for various types of crops, many studies have shown a positive relationship between VH backscatter coefficient in C-band and LAI or NDVI (Mandal et al., 2019; Wang et al., 2019), our results show that, in deciduous forests, these relationships are inverted. Temporal dynamics of LAI and NDVI is accompanied by a decrease in VH and an increase of VV/VH ratio. These relationships are also not unique but depend on the phenological stage considered. The use of S-1 signal in classifications or in fusion with optical remote sensing or its interpretation should be considered with great care.

## 5. Conclusion

Time-series of Sentinel-1 A and B backscattering coefficients were used to characterize the seasonal phenological cycles in a temperate deciduous forest over five years. While the backscattering coefficient in vertical polarization (σ^0^VV) remains relatively stable over the seasons, the backscattering coefficient in cross-polarization (σ^0^VH) responds significantly to the seasonal vegetation cycle, with a behavior that is opposite to what is usually observed on various crops. σ^0^VH decreases during spring simultaneously with spring budburst and leaf expansion, reaches a minimum during the main growing season when canopy leaf area is at its maximum and increases again simultaneously with leaf fall. The observed σ^0^VH seasonal amplitude is 1.8 dB on average. S-1 time series of VV/VH ratio provides a good description of the seasonal vegetation cycle, allowing the extraction of spring and autumn phenological markers. Estimates of budburst dates in spring differ by approximately 8 days on average from field phenological observations. During the senescence phase, the estimates provided by VV/VH are late by about 20 days in comparison to field phenological observations and deviate significantly by about two to four weeks from the estimates provided by *in situ* NDVI, S2-based NDVI and LAI time-series. While during the spring, temporal pattern of VV/VH correlates well with LAI and NDVI, during senescence, it is better explained by the dynamics of litterfall. The deviation between VV/VH and litterfall-based senescence estimates is reduced to about one week. A hysteresis phenomenon is observed on the relationships between VV/VH and NDVI or LAI. For the same LAI or NDVI, the response of VV/VH is lower during canopy foliage increase in spring than during leaf senescence and fall in autumn. These relationships are not unique and lead to the conclusion that the mechanisms involved in the seasonality of VV/VH signal are different according to the phenological stage considered. This behavior can be explained by the preponderant contribution of woody parts in VH polarization backscattering, which decreases as the forest canopy becomes more and more closed during spring and summer and increases again during leaf senescence and fall in autumn. The hysteresis phenomenon shows that attenuation of σ^0^VH signal by canopy foliage appears to be less important during spring and early summer than during senescence. This may be caused by two opposing mechanisms: a significant role of water content of canopy foliage during the spring which has the effect of causing an increase of σ^0^VH and the role of foliage, which masks the woody components, and has an opposite effect of causing a decrease in σ^0^VH. This last mechanism appears to be preponderant since σ^0^VH decreases during the spring transition as shown in this study but also in previous studies. During the senescence, the strong relationship between the σ^0^VH increase and the opening of the canopy caused by litterfall suggests that the masking role played by foliage is also dominant. These results show that the interpretation of S-1 signals over deciduous forest canopies or their use for classification without or after fusion with optical data must be carried out with great care due to the temporal variability of the contributions of the different canopy components associated with the seasonal phenological cycle. The use of physical approaches based on radar backscatter models in forest canopies will have the advantage of allowing a better understanding and evaluation of the contributions of the different ecosystem components in the measured signal. It will also provide useful information to better establish the correspondence between indirect phenological metrics predicted from the S-1 time series and field phenological observations.

## Supplementary Materials

**Figure S1:**
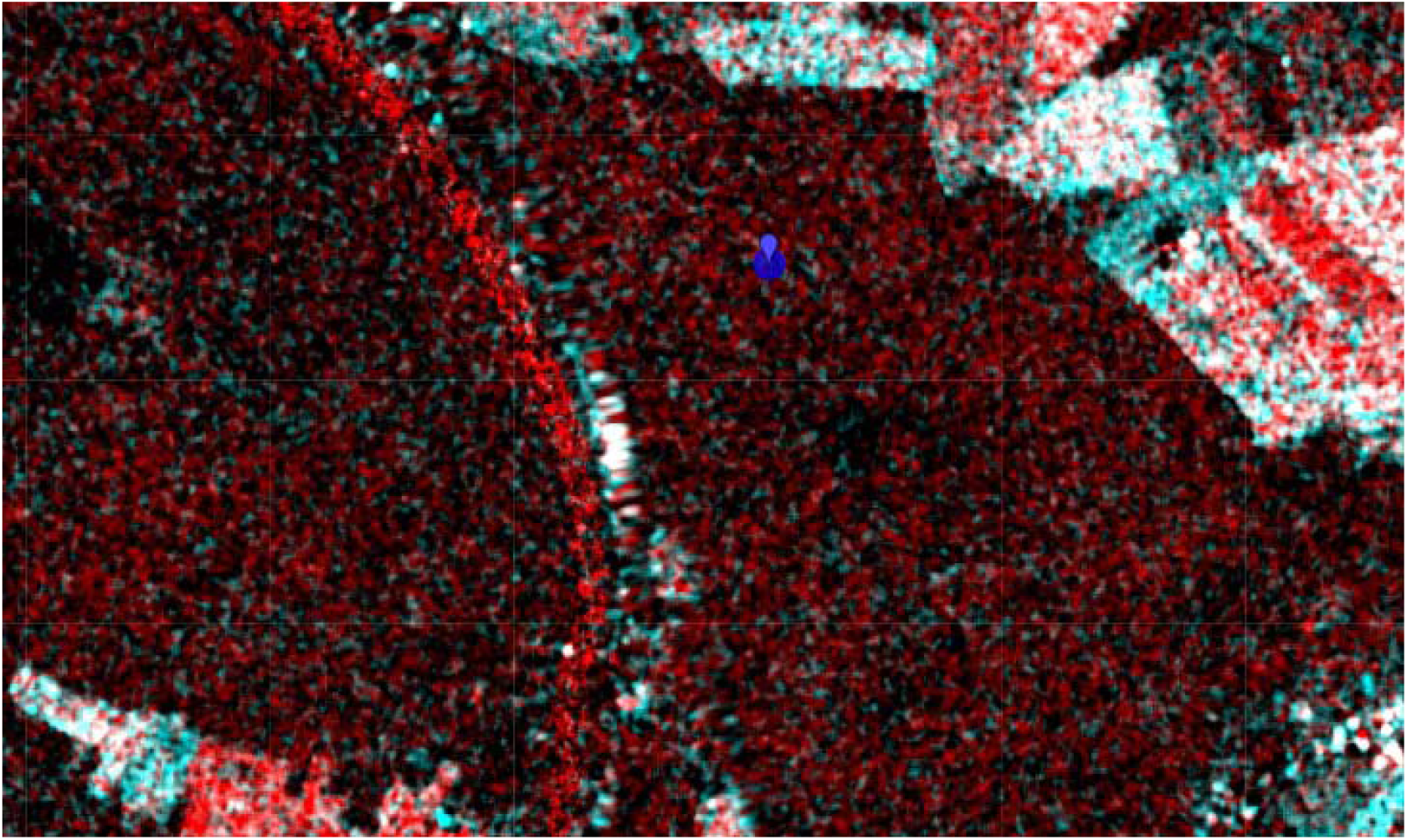
Fontainebleau-Barbeau forest location in France (48°28’26”N, 2°46’57”E). The blue circle with a radius of 50 m is centered on Fontainebleau-Barbeau flux tower (FR-Fon, ICOS network). The image is a Sentinel-1 RGB composite where average VV/VH during the summer (15/06/2018 - 31/08 2018) in red, average VV/VH during the winter (01/01 – 28/02 2018) in green and blue.

**Figure S2:**
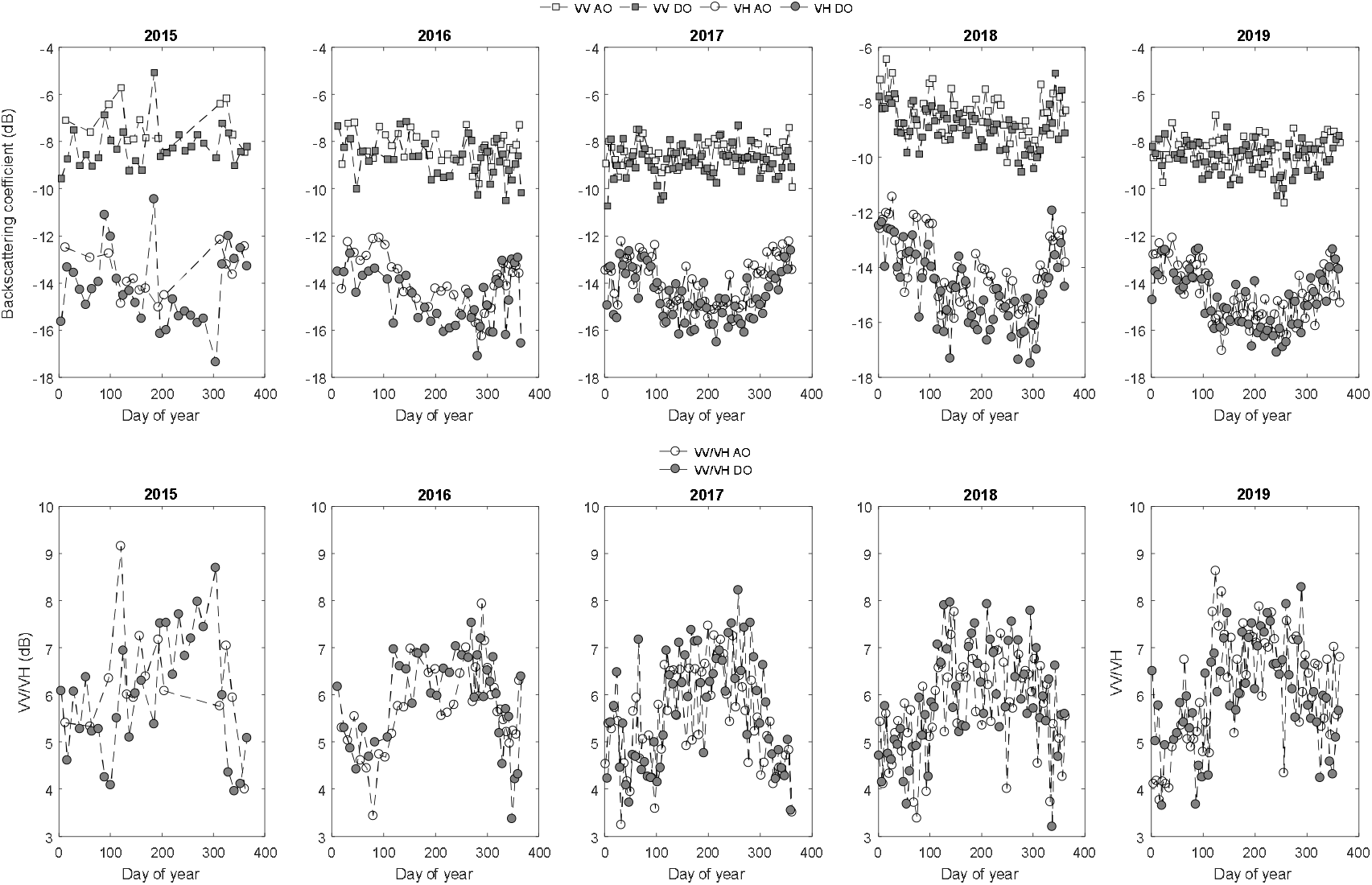
SAR backscattering coefficients σ^0^ in VV and VH polarizations and VV/VH ratio time-series from Sentinel-1 A&B. circle: VH polarization, square: VV polarization. Empty (circles and squares) in ascending orbit and filled in descending orbit.

**Figure S3:**
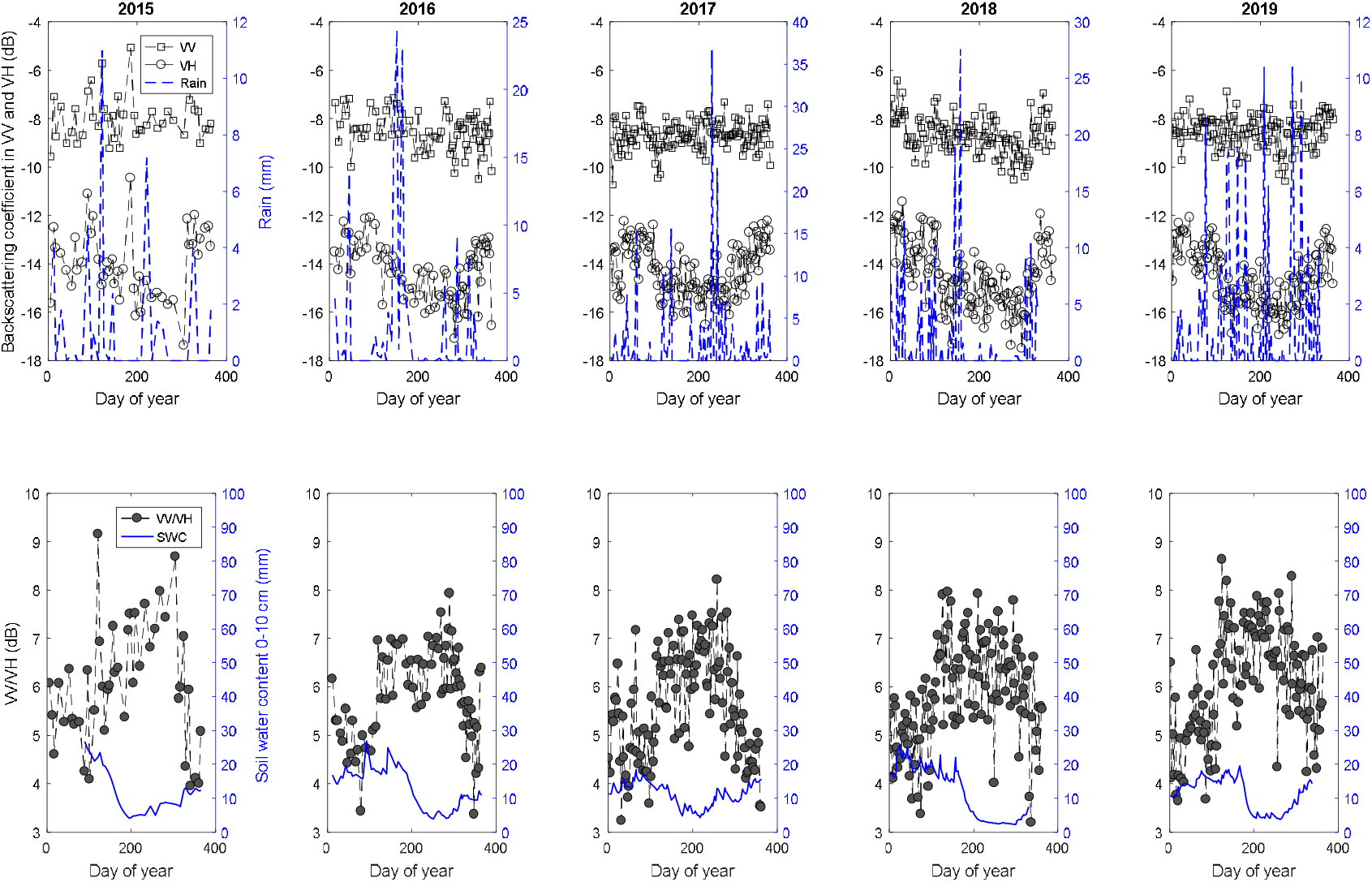
Sentinel-1 σ^0^VV (square symbol), σ^0^VH (circle symbol) and VV/VH (filled circle) time-series over 2015-2019 period (right axis). In blue (left axis), rain (upper panel) and soil water content (0-10 cm) in mm (lower panel).

**Figure S4:**
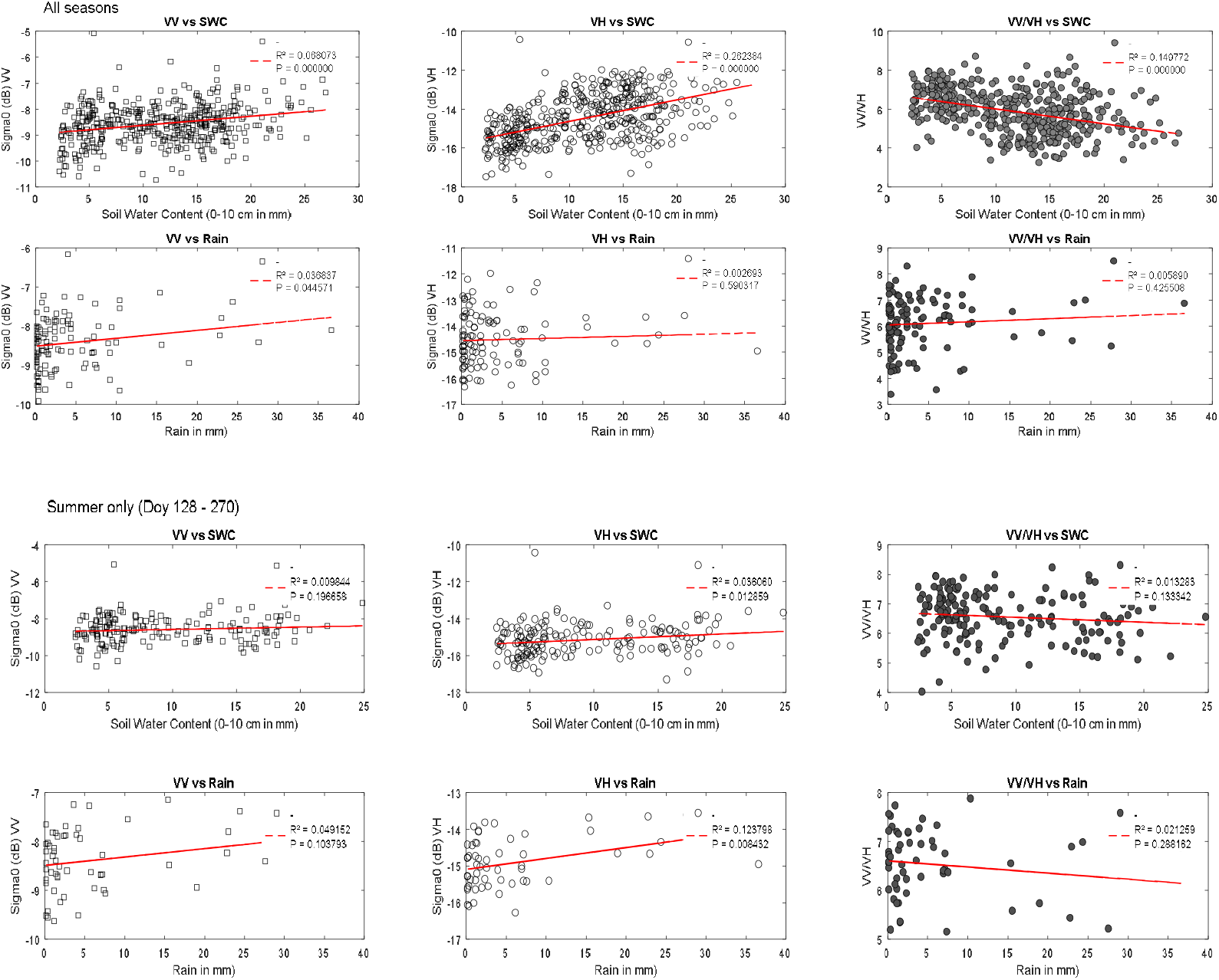
Linear regressions between Sentinel-1 data and soil water content (0-10 cm depth) and rain during the year (all seasons – upper panel) and during summer season (lower panel).

**Table S5:** observed and estimated dates of onset of greenness in spring and leaf senescence and fall based on field phenological observations and time-series of Sentinel-1 VV/VH, ground-based normalized difference vegetation index (NDVI), Leaf area index (LAI), Litterfall and Sentinel-2 NDVI.

|  | Budburst and leaf expansion |  |  | Leaf senescence and fall |  |  |
| --- | --- | --- | --- | --- | --- | --- |
|  | SOS | MOS | EOS | SOE | MOE | EOE |
| Phenological observations |  |  |  |  |  |  |
| 2015 | 101 | 106 | 109 | 267 | 290 | 312 |
| 2016 | 102 | 109 | 115 | 282 | 300 | 317 |
| 2017 | 89 | 99 | 108 | 267 | 294 | 321 |
| 2018 | - | 105 | - | - | - | - |
| 2019 | - | 109 | - | - | - | - |
| Sentinel-1 VV/VH |  |  |  |  |  |  |
| 2015 | 110 | 113 | 113 | 312 | 323 | 332 |
| 2016 | 100 | 123 | 144 | 304 | 321 | 336 |
| 2017 | 100 | 117 | 130 | 273 | 297 | 318 |
| 2018 | 89 | 104 | 115 | 292 | 323 | 350 |
| 2019 | 99 | 111 | 121 | 252 | 321 | 388 |
| In situ NDVI |  |  |  |  |  |  |
| 2015 | 102 | 107 | 112 | 272 | 293 | 314 |
| 2016 | 103 | 113 | 123 | 280 | 302 | 324 |
| 2017 | 90 | 101 | 111 | 255 | 293 | 332 |
| 2018 | 100 | 106 | 112 | 272 | 311 | 349 |
| 2019 | 96 | 108 | 119 | 273 | 306 | 339 |
| Leaf Area Index |  |  |  |  |  |  |
| 2015 | 100 | 113 | 124 | 251 | 287 | 322 |
| 2016 | 103 | 120 | 137 | 263 | 296 | 329 |
| 2017 | 90 | 111 | 131 | 248 | 288 | 327 |
| 2018 | 102 | 111 | 119 | 231 | 287 | 341 |
| 2019 | 98 | 115 | 131 | 250 | 292 | 334 |
| Litter fall |  |  |  |  |  |  |
| 2015 | - | - | - | 266 | 301 | 309 |
| 2016 | - | - | - | 298 | 312 | 324 |
| 2017 | - | - | - | 273 | 300 | 322 |
| 2018 | - | - | - | 281 | 316 | 331 |
| 2019 | - | - | - | 288 | 317 | 329 |
| S-2 NDVI |  |  |  |  |  |  |
| 2015 |  |  |  |  |  |  |
| 2016 | 115 | 118 | 124 | 261 | 285 | 320 |
| 2017 | 105 | 113 | 121 | 303 | 316 | 328 |
| 2018 | 106 | 111 | 115 | 192 | 281 | 368 |
| 2019 | 83 | 99 | 114 | 253 | 296 | 337 |

